# Rapid detection of identity-by-descent tracts for mega-scale datasets

**DOI:** 10.1101/749507

**Authors:** Ruhollah Shemirani, Gillian M. Belbin, Christy L. Avery, Eimear E. Kenny, Christopher R. Gignoux, José Luis Ambite

**Affiliations:** Information Sciences Institute, University of Southern California, Marina del Rey, CA; Center for Genomic Health, Icahn School of Medicine at Mount Sinai, New York, NY; The Charles Bronfman Institute of Personalized Medicine, Icahn School of Medicine at Mount Sinai, New York, NY; Department of Epidemiology, University of North Carolina at Chapel Hill, Chapel Hill, NC; Department of Genetics and Genomic Sciences, Icahn School of Medicine at Mount Sinai, New York, NY; Department of Medicine, Icahn School of Medicine at Mount Sinai, New York, NY; Colorado Center for Personalized Medicine, University of Colorado Anschutz Medical Campus, Aurora, CO; Department of Biostatistics and Informatics, University of Colorado Anschutz Medical Campus, Aurora, CO

## Abstract

The ability to identify segments of genomes identical-by-descent (IBD) is a part of standard workflows in both statistical and population genetics. However, traditional methods for finding local IBD across all pairs of individuals scale poorly leading to a lack of adoption in very large-scale datasets. Here, we present iLASH, IBD by LocAlity-Sensitive Hashing, an algorithm based on similarity detection techniques that shows equal or improved accuracy in simulations compared to the current leading method and speeds up analysis by several orders of magnitude on genomic datasets, making IBD estimation tractable for hundreds of thousands to millions of individuals. We applied iLASH to the Population Architecture using Genomics and Epidemiology (PAGE) dataset of ∼52,000 multi-ethnic participants, including several founder populations with elevated IBD sharing, which identified IBD segments on a single machine in an hour (∼3 minutes per chromosome compared to over 6 days per chromosome for a state-of-the-art algorithm). iLASH is able to efficiently estimate IBD tracts in very large-scale datasets, as demonstrated via IBD estimation across the entire UK Biobank (∼500,000 individuals), detecting nearly 13 billion pairwise IBD tracts shared between ∼11% of participants. In summary, iLASH enables fast and accurate detection of IBD, an upstream step in applications of IBD for population genetics and trait mapping.

---

Inferring segments of the genome inherited Identical-By-Descent (IBD) is a standard method in modern genomics pipelines to understand population structure and infer relatedness across datasets^1-6^. Furthermore, it can be leveraged for alternative mapping strategies such as population-based linkage^7^, capturing rare variation from array datasets^8^, and improving long-range phasing. However, the ability to scale this process to mega-scale datasets while comparing individuals along the genome has been limited. While approximate methods have been developed to improve phasing^9-11^, the identification of accurate segments inherited IBD has been limited, making its integration with variant-based testing challenging in the modern genomic context. Here we extend ideas originally applied to large-scale similarity detection^12^ to develop iLASH, IBD by LocAlity-Sensitive Hashing, a novel algorithm that provides ultra-rapid and sensitive computation of identity-by-descent. In contrast to previous methods, which are based on conventional hashing techniques (e.g., GERMLINE^13^), iLASH uses a locality sensitive hashing (LSH) technique on short slices of DNA array data of individuals so that DNA slices near each other are located on an IBD tract with high probability, while data points far away from each other are likely to be on different IBD tracts. We provide further speedups via a careful implementation that leverages multiple processing cores, now commonly available in modern CPUs, through parallelizing computation in multiple stages of the algorithm. This parallelization also takes advantage of idle cycles during file reading and writing.

LSH algorithms are a category of hashing functions that preserve distances while maintaining computational efficiency in high dimensional applications^14^. LSH algorithms have been shown to be efficient in machine learning^15^, entity linkage^16^, search engines^17,18^, and other disciplines^19,15^. Here, we introduce a modification of the LSH algorithm designed expressly for identity-by-descent detection on genomic data. Based on the minimum length of tracts being searched for, iLASH slices the haplotypic data of every individual into windows and then performs LSH separately on each slice. After identifying slices that have a high probability of being in contiguous IBD, iLASH extends the slices efficiently to find the precise borders of each IBD tract. Multiple measures have been put in place to speed up the estimation process, including native parallelization and adaptive hashing. To maximize efficiency, iLASH also uses idle input time to preprocess the loaded genotype data. Similarly, it uses idle output time for calculating the extended IBD tracts and performing calculations of length and exact similarity. Furthermore, iLASH precomputes the permutations needed for LSH, a simple step that enables the threads to independently calculate LSH signatures in parallel. Overall, a combination of algorithmic technique and low-level optimizations allows for increased efficiency in large-scale genomic investigations.

The framework of the iLASH algorithm is described next and is shown in Figure 1 with additional details available in the online Methods section. The algorithm relies on phased haplotypes from a typical GWAS array with hundreds of thousands to millions of variants (SNPs) represented as binary strings. iLASH builds upon a form of hashing originally developed to detect similar entries in large document databases. We are only interested in the similarity of segments longer than a given threshold, as the probability of a shared haplotype being IBD increases as segments grow longer^13,20^. Therefore, the first step of iLASH is to break the genetic data of the population into slices of a prespecified genetic length around the threshold of interest, say 3cM (Figure 1). Each slice can be processed in parallel until the last step of the algorithm. Second, we break each slice into segments of *k* bits (aka k-mers, or “shingles”, with k typically 10-30 bits), either in a contiguous or overlapping fashion. The genetic data for all the individuals in a slice is then transformed into sets whose elements are the distinct k-mers. Third, we compute the *minhash signature*^16^ of these sets as illustrated in the second panel of Figure 1^21^. The *minhash* approach provides a very efficient way of approximating the Jaccard similarity between the sets using a sublinear number of comparisons, efficiently scaling in large datasets. To create the *minhash* signature matrix, iLASH generates random permutations of the k-mers in a slice, and for each permutation it records the first index at which an individual’s shingle set at that slice contains a value, called the *minhash* value of the individual for that permutation at that slice. For example, in Figure 1 for individual I3 and permutation P2 (H3, H5, H1, H4, H2), the first element in the k-mer set I3 following the order of permutation P2 that has a non-empty value is H1, which is the 3rd element of the permutation, thus the minhash(I3,P2)=3. The probability of two individuals having the same *minhash* value is equal to the Jaccard similarity of their k-mer sets. Hence, the Jaccard similarity can be estimated by generating *minhash* values using different permutations and comparing them; with the estimation error decreasing as the number of permutations increases. For example, even with the low number of 3 permutations in Figure 1, the intersection of *minhash* signatures and the Jaccard similarity of I1 and I3 coincide. However, it would be computationally inefficient to compare all pairs of signatures. The banding technique used to compute LSH signatures in the next step allows for significant speedup by only selecting candidates for comparison that have an increased probability of having a high Jaccard similarity and avoiding most of the possible comparisons. We group the *minhash* signatures into b bands comprised of r *minhash* values and hash each band. Assume that the Jaccard similarity of I1 and I2 is s. It can be shown^21^ that the probability that two individuals agree in at least one LSH hash value is 1 - (1 - s^r^)^b^. This function is logistic with a step transition controlled by parameters r and b, which can be tuned to trade-off accuracy of approximation versus efficiency. Finally, iLASH uses these similar LSH hashes to find candidate IBD matches and then examines neighboring slices of a match candidate pair to extend them to identify the full IBD segment (Figure 1).

**Figure 1.**
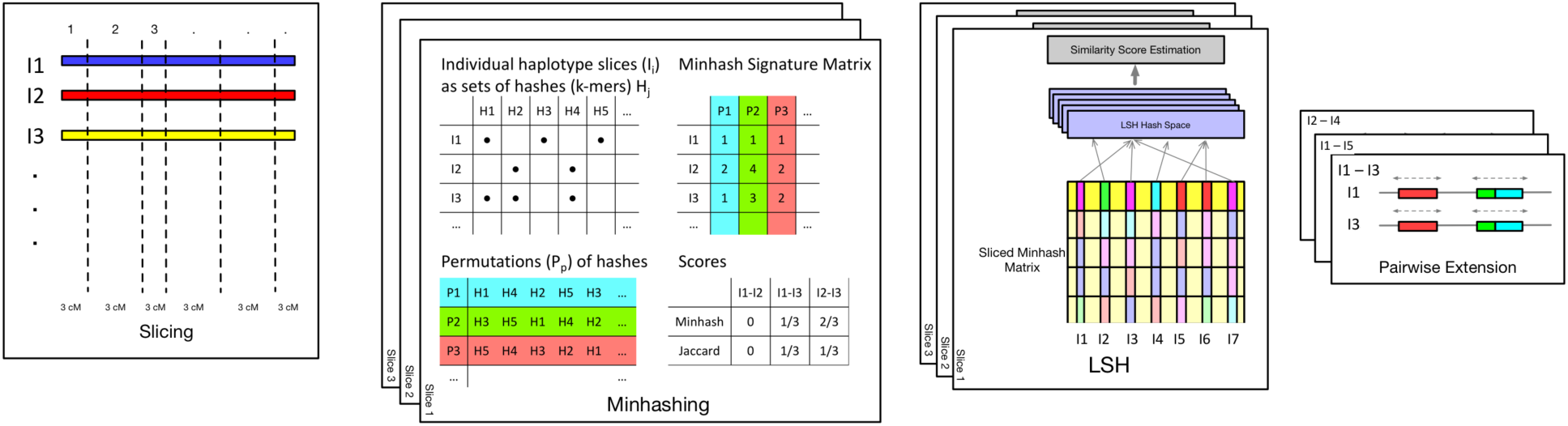
Schematic of the iLASH algorithm pipeline. Starting from the left with the ***Slicing*** step where haplotypes are broken into windows (of uniform or variable length). The ***Minhashing*** step creates minhash signatures by generating a table of random permutations. The ***LSH*** step bands together minhash values to create an integrated LSH hash table where candidate matches are grouped together. Finally, in the ***Pairwise Extension*** step, these candidates are further analyzed to be extended in the (likely) case that an IBD tract spans multiple windows.

iLASH takes standard genotype files in plink^22^ format as input and is available open source. For efficiency purposes, iLASH is implemented in C++. To foster usability, iLASH is designed to run on a single machine with one command with sensible defaults derived from realistic simulations. However, it is highly configurable to allow users to tune its behavior for each dataset, e.g., arrays of different densities.

## Results

We present a thorough evaluation of iLASH performance for both simulated data and real data from the PAGE consortium and the UK BioBank. The size of our test datasets is intractable for common tools, such as PLINK and Beagle, so we do not compare with them. We compare iLASH to the standard algorithm, GERMLINE^13^, for both performance and accuracy. Other than being the first tool with efficient performance for comparison to iLASH, GERMLINE is also similar to iLASH in that it uses hashing methods for IBD inference. We also separately compare iLASH against RaPID^23^, a recently developed scalable IBD algorithm, for both performance and accuracy.

### Performance on Simulated Data

To investigate iLASH performance, we simulated IBD haplotypes for three separate populations with different average IBD and for sizes ranging from 1,000 to 80,000 individuals. To create these data, we first used HapGen^24^ to simulate 80,000 individuals with an elevated error rate in the copying model (the error rate for a given SNP (Θ) =130) to decrease background IBD. Then we scanned three populations in the PAGE dataset with different cryptic relatedness characteristics: African American (low IBD on average), Puerto-Rican (a founder population with elevated IBD sharing), and all the individuals in PAGE, and extracted their detailed IBD distributions. We used these IBD distributions to generate “ground truth” IBD segments set with the same number of segments and lengths observed in the mentioned populations among any randomly selected group of 1,000 samples. We repeated this process to create a ground truth IBD map for 80,000 samples. The Puerto Rican population IBD simulation, for example, has more than 10 million shared segments with a total length of 62 million cM.

#### Accuracy

Using our simulated data as ‘ground truth’ we compared the accuracy of iLASH & GERMLINE. iLASH accurately recovers at least 95% of all simulated IBD haplotypes across the simulated dataset of the three populations tested, compared to a lower accuracy of 82% for the GERMLINE algorithm, results that were consistent when considering 1,000 to 30,000 individuals for both GERMLINE and iLASH (see Supplemental Figure 1). However, the concordance with ground truth varies significantly with the tract length, as shown in Figure 2, with iLASH outperforming GERMLINE both on shorter tracts and on longer tracts when high accuracy is required. For example, for tracts of 3cM, iLASH identifies approximately 80% of tracts with at least 90% accuracy, while GERMLINE only identifies 35% of those tracts. For tract lengths of 5, 10, or 20cM, iLASH has recall, or the proportion of true tracts identified, close to 100% for accuracies up to 95%.

**Figure 2.**
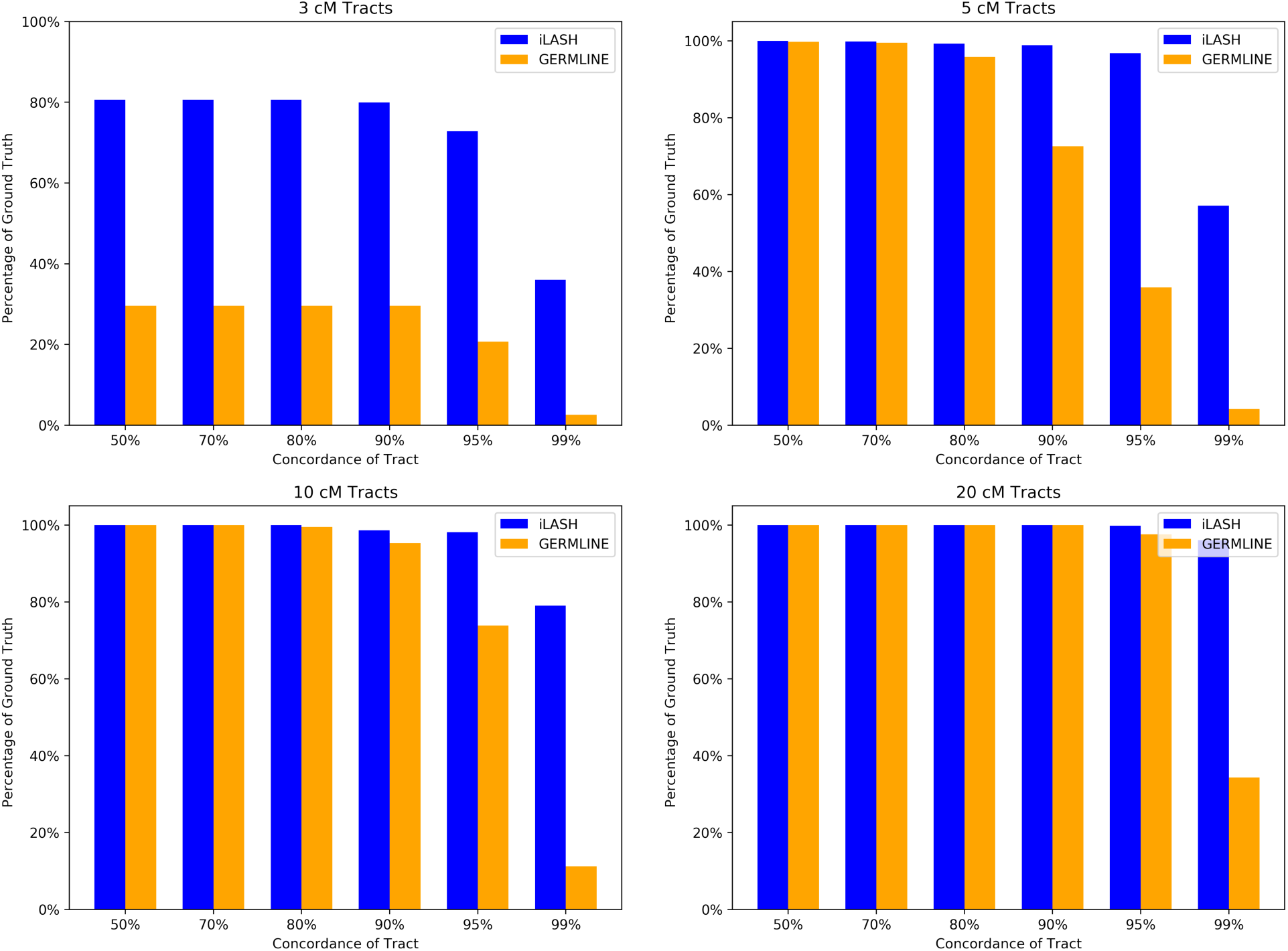
Concordance of iLASH and GERMLINE with ground truth on simulated data at tract lengths of 3, 5, 10, and 20cM and accuracies from 50% to 99%.

#### False Positive Rate

To investigate the rate of false positives of iLASH and GERMLINE, we took a composite individuals approach^25^. We used chromosome 2 data from 2000 sampled individuals from the African American population of the PAGE study. In order to minimize the possibility of false positive IBD inference, we divided the genotype data into slices of 0.2 cM length, and we shuffled theses slices among the 2000 individuals. Before this process, iLASH was able to find 10459 IBD tracts in the dataset. Afterwards, however, only 1 tract of 3.9 cM was found by iLASH. In the same reshuffled dataset, GERMLINE still found 27,838 (false positive) tracts, with 99.9% of them around a low complexity region (starting from SNP rs1729898 and ending at SNP rs12464090, which contains only 60 SNPs on the PAGE array data). After trimming the genotype data by dropping one out of every 3 SNPs to see how iLASH performs in the context of less density, the number of tracts found by iLASH decreased to zero. GERMLINE results also decreased to 1,097, all of which were still in the same region. When we increased the size of slices to 0.5 cM, iLASH identifies 3 tracts for the normal data and 13 tracts for the data with lower SNP density. The number of GERMLINE results were 22,688 and 26,724 tracts for dense and trimmed haploid files respectively. Again, more than 99% of the false positive tracts found by GERMLINE were located in the same low complexity region described above. In contrast, iLASH showed near perfect performance in both cases. It is worth noting that since both tools use haplotype data instead of genotype data, their precision is dependent on the accuracy of the phasing stage, however standard phasing algorithms embrace approximate IBD inference to improve long-range phasing. Such methods are expected to improve phasing accuracy in large studies, particularly in the haplotypes spanning IBD segments.

#### Runtime

To compare the time efficiency of iLASH and GERMLINE, we used the same simulated datasets as the previous section. We ran GERMLINE and iLASH on the same machine, a workstation running CentOS Linux release 7.4.1708 with 128 GB of memory and Intel® Xeon® Processors E5-2695 v2 with 12 cores and 24 threads on it using 30 MB of shared cache. Both iLASH and GERMLINE are implemented in C++, but in order to use a specific version of the Boost library, GERMLINE required an earlier version of the C++ compiler.

Figure 3(A) shows the computation time in seconds for iLASH and GERMLINE as the population size grows. The biggest dataset that we were able to run GERMLINE tractably on one machine was for 30,000 individuals and took over 5 hours to run. For the same data, iLASH took 3 minutes and 15 seconds. Our maximum-sized simulation of 80,000 individuals could be scanned and analyzed by iLASH in less than 16 minutes.

**Figure 3.**
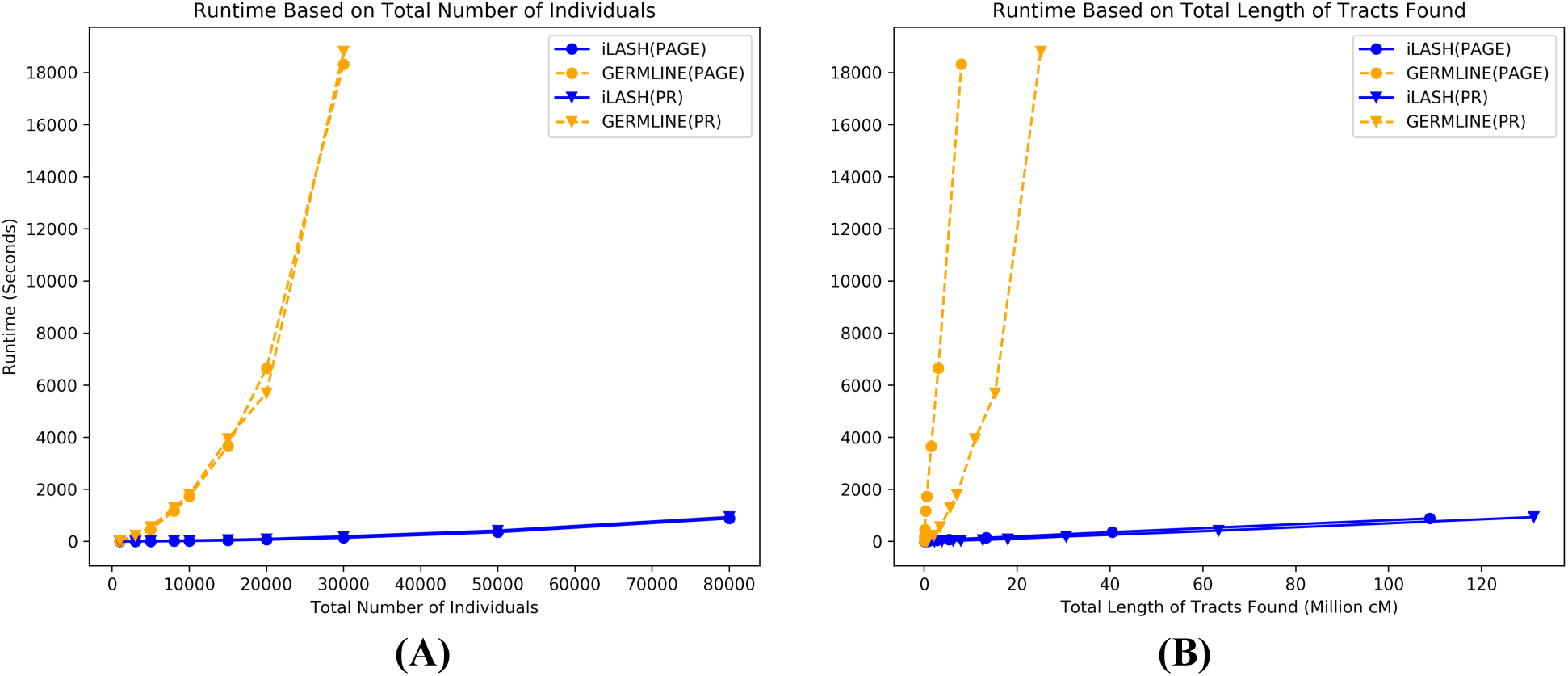
IBD computation runtime in seconds for iLASH and GERMLINE on synthesized haplotypic data simulating all of PAGE and Puerto Rican (PR) populations IBD patterns: **(A)** as the number of individuals grows, **(B)** as the total output (total length of tracts found) grows.

Figure 3(B) shows the computation time in seconds for iLASH and GERMLINE as the total size of found IBD grows. Both tools exhibit a linear growth with the amount of IBD, but iLASH’s slope if significantly lower than GERMLINE’s (time_iLASH_ = 0.8 x IBDLength vs. time_GERMLINE_ = 229 x IBDLength; fitted over the PAGE dataset).

### Comparison with RaPID

Recently, Naseri et al.^23^ released an IBD detection package that relies on the Positional Burrows-Wheeler Transfom^26^ to efficiently scale haplotype matching. We ran RaPID on the same simulated data used in our comparisons with GERMLINE and iLASH. We noted that while RaPID is substantially faster than GERMLINE, it remains slower than iLASH on all sized datasets, and the difference is particularly noticeable as the sample size increases (cf. Supplemental Figure 2). For 1,000 samples, iLASH takes 4 seconds, while RaPID takes 27 seconds. For 80,000 samples iLASH takes 15 minutes, while RaPID takes 98 minutes. More importantly, iLASH is significantly more accurate than RaPID: iLASH recovered over 95% of the total length of “ground truth” IBD in simulations, where RaPID only recovered 72%. Short IBD segments (5-3cM) were particularly challenging for RaPID, which generated a larger number of false positives (22-25% across runs). In spite of RaPID being a haploid method, the output of the program does not report haplotype phase information, which somewhat constrains the options possible in downstream analysis after IBD estimation. Given these limitations, in the remainder of the paper, we only compare with GERMLINE.

### Performance on real data from the PAGE Study

We investigated iLASH and GERMLINE IBD inference over two existing datasets: a multi-ethnic population of N=51,520 individuals from the Population Architecture using Genomics and Epidemiology (PAGE) consortium, and the N**∼**500,000 individuals in the UK Biobank dataset. In PAGE, iLASH uncovered a total 202,424,985 segments, while GERMLINE identified a total of 195,577,460 tracts. The overall concordance between iLASH and GERMLINE was 95%. iLASH total runtime was 58 minutes on a single workstation (same as above) requiring between 3 GB (for chromosome 22) and 17 GB of memory (for chromosome 2). GERMLINE could not be run on the same workstation, because it required more than 128 GB of memory for every chromosome. GERMLINE was run on the High Performance Computing Cluster called Minerva at the Icahn School of Medicine at Mount Sinai, which has several high-memory nodes. For the largest chromosome (12) that could be analyzed by GERMLINE without splitting the population, GERMLINE took 6 days of computation. For the same chromosome in the single machine described above, iLASH took 3 minutes and 12 seconds, an improvement of four orders of magnitude.

To explore the utility of IBD haplotypes inferred by iLASH in a large genomic dataset we constructed an IBD-based network of distant relatedness among the PAGE dataset^27^. In this network individuals are represented by nodes (N=38,919 across 3 PAGE Studies: WHI, MEC and HCHS/SOL) that are connected by edges (e= 55,069,482) if they share any haplotypes IBD. We used this to explore fine-scale population substructure by applying the community detection algorithm InfoMap^28^ to the IBD network in order to uncover communities of individuals who were enriched for recent, shared ancestry in the form of elevated IBD sharing. We observed that 92.3% of selected PAGE participants fell into one of 12 inferred IBD communities, each containing N>100 individuals, with the remaining 7.7% of participants being assigned to communities ranging from N=1 to N=91 participants in size (see Figure 4A). We observed that IBD community membership correlated strongly with available demographic information in the form of both selfreported ethnicity as well as sub-continental and country level region of origin (see Supplementary Table 1). For example, membership of one InfoMap community was highly correlated with being born in Puerto Rico (PPV 0.96), while another was correlated with being born in the Dominican Republic (PPV 0.98). We also observed significant differences in the distribution of total pairwise IBD sharing between communities (Figure 4B). Examination of the population level fraction of IBD sharing within- and between-communities revealed a high degree of network modularity, or elevated sharing within communities relative to between (Figure 4C). Three distinct communities emerged that correlated with being born in Mexico (PPVs 0.96, 0.43 and 0.99, respectively), the latter of which exhibited elevated IBD sharing relative to the former two and may represent a founder population of (native American) Mexican origin. This analysis demonstrates the utility of IBD inference for exploring fine-scale population substructure within large datasets. Further, this elevated IBD signature empowers techniques in founder populations such as IBD mapping and detection of highly drifted alleles.

**Figure 4.**
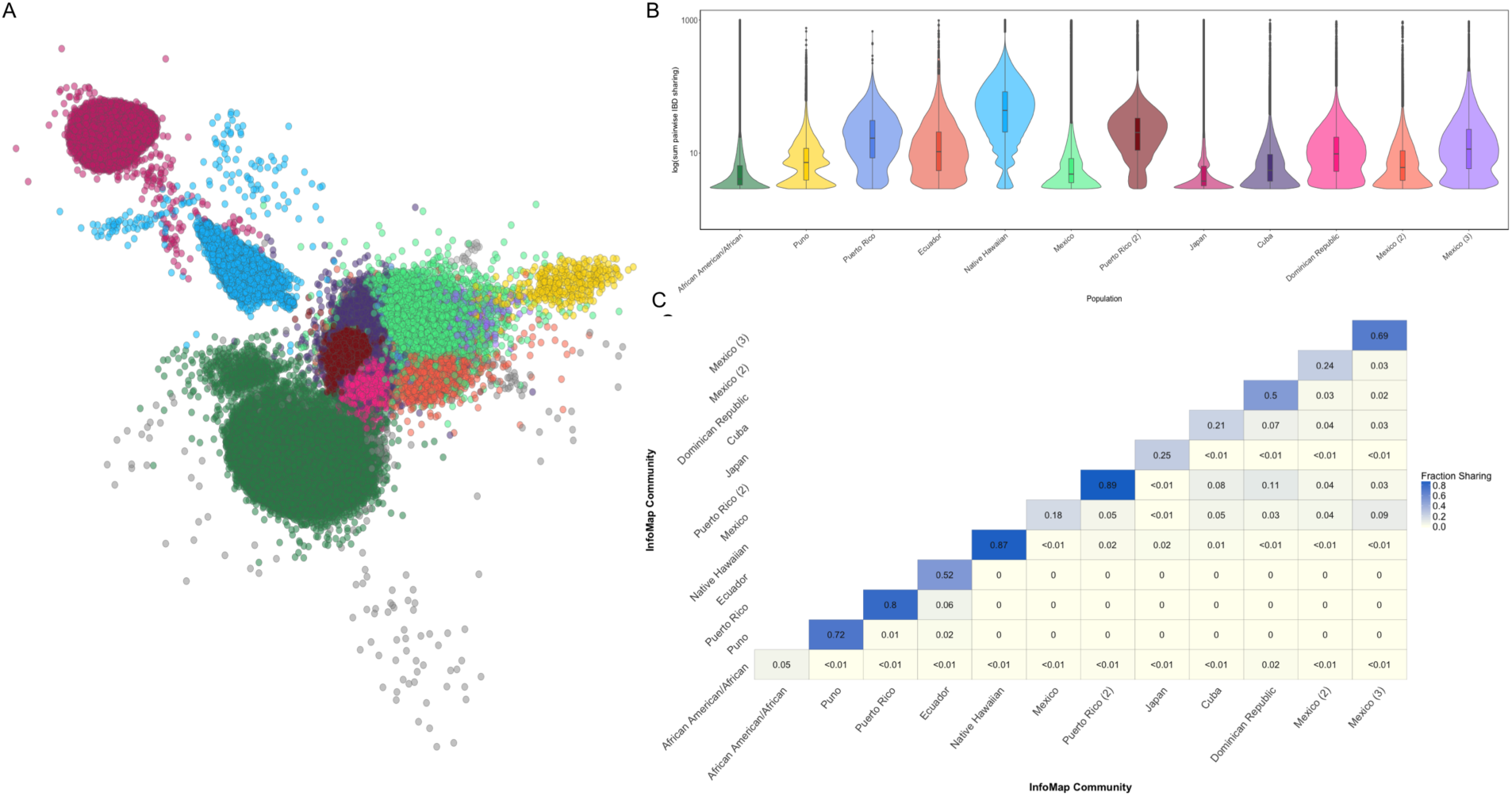
Network of IBD sharing in the PAGE dataset. (**A**) A network of IBD sharing within PAGE plotted via the Fruchterman Reingold algorithm. Each node represents an individual (edges not shown). Individuals are colored based on community membership as inferred by the InfoMap algorithm. (**B**) Distribution of the sum of IBD sharing within the top 16 largest InfoMap communities demonstrates variance in levels of IBD sharing between different communities. InfoMap communities are labeled according to the demographic label that most strongly correlated with community membership (as measured by positive predictive value). Elevated pairwise IBD sharing can be observed in several InfoMap communities, which may represent founder effects. (**C**) Heatmap of the population level fraction of IBD sharing within and between the top 16 largest InfoMap communities demonstrates elevated sharing within, relative to between communities.

### Detecting Identity-by-Descent in the UK Biobank

To explore fine-scale population substructure in the UK Biobank we leveraged the phased genotype data at 655,532 SNPs for N=487,330 participants. We used iLASH to infer pairwise IBD segments (>=2.9cM) between all individuals. We observed 10.84% of all possible pairs of individuals shared at least one haplotype of their genome IBD, representing 12,867,760,228 pairwise connections in total (Figure 5A). To understand how well the IBD sharing estimates correlated with genetic relatedness, we calculated the correlation between the kinship coefficient and the sum of IBD haplotype sharing among 3rd degree and above relationships in the UK Biobank. We observed a high degree of correlation between the two estimates (R^2^=0.95; Figure 5B). Beyond this close relatedness, we observed 778,822 pairs of individuals exhibiting relatively high levels of relatedness (>100cM), and additionally 43,205,248 pairs of individuals with sharing above 20cM. In most instances these individuals were just below the level of detection as provided by the standard genetic relationship matrix released alongside the UK Biobank genotype data. However, we also identified 4,808 highly related pairs of individuals (>=500 cM) that were not reported to be 3^rd^ degree relatives or above in the default UK Biobank release. To investigate this further, we replicated the KING relatedness estimation for this subset of participants, and noted that the majority of these pairs did exhibit elevated relatedness (mean kinship=0.037, Interquartile Range=0.031-0.043), but that their KING estimates fell slightly below the cut-off for 3^rd^ degree relatives (>0.0442). However, some discordance between the two metrics persisted. Specifically, we identified a subset of pairs (N=203 pairs, comprised of N=378 unique individuals) with high IBD (>500cM), but low or negative kinship (< 0.02). We noted that levels of autozygosity within the individuals comprising these pairs was significantly elevated relative to the population average in the UK Biobank, with the mean sum of runs of homozygosity (ROH) within discordant pairs being 116.5cM (95% C.I=98.2-135.0cM), compared to 1.84cM (95% C.I=1.79-1.89cM, Wilcoxon p<6.3e-204) in the UK Biobank overall. We speculate that this elevation of autozygosity may have contributed to the deflation of the KING kinship estimates and resultant discordance with iLASH.

**Figure 5.**
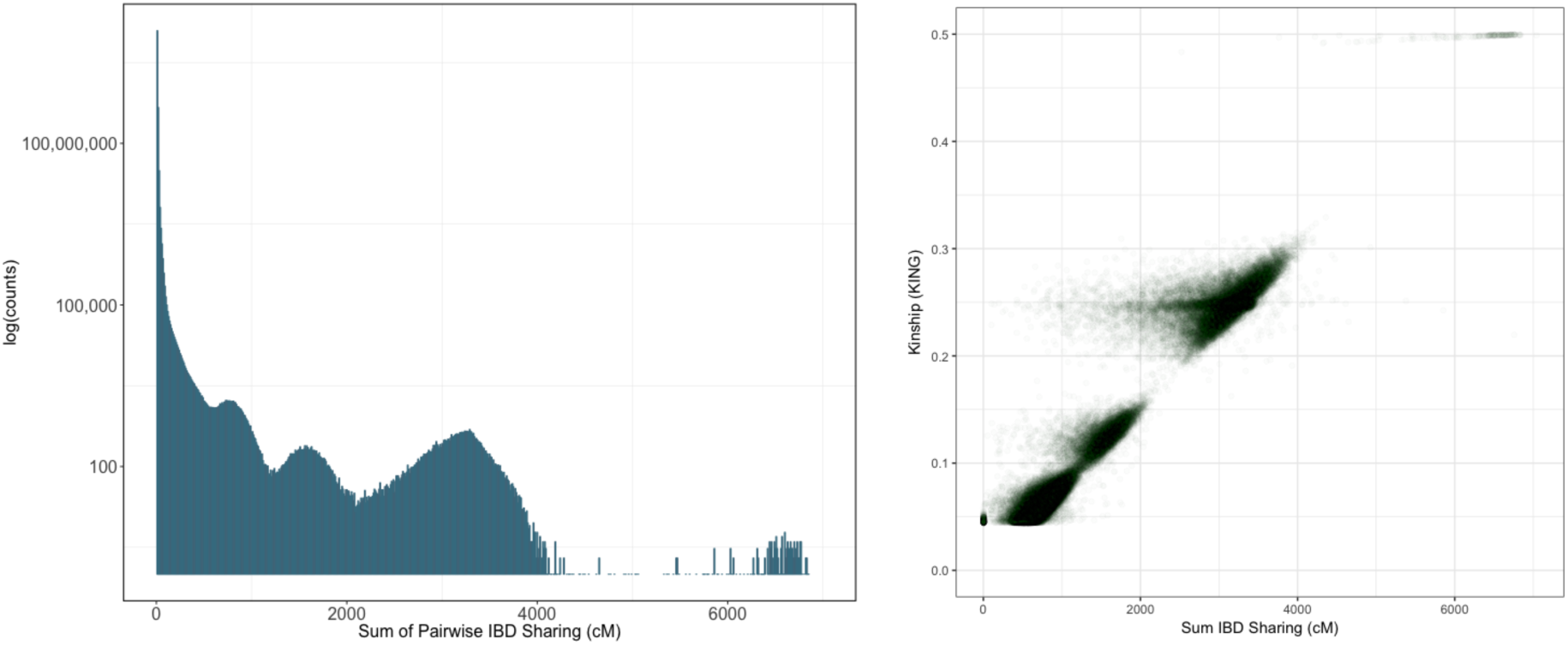
Identity-by-Descent Sharing in the UK Biobank. (**A**) Distribution of the sum of pairwise IBD sharing (cM) in the UK Biobank across all N=487,330 participants. (**B**) Correlation between the sum of IBD sharing and kinship as measured by the KING software in all pairs of individuals reported in the UK Biobank output to be >= 3^rd^ degree relatives.

Overall, we highlight in the UK Biobank that detectable relatedness exists across far more pairs of individuals than is present in the kinship matrix currently provided with the standard data release. The sensitivity of methods like iLASH to detect true levels of cryptic relatedness is critical in mixed effects models such as SAIGE^29^ and BOLT-LMM^30^ for efficient, appropriately-calibrated association testing.

## Discussion

Here we present iLASH, an accurate and computationally efficient method for detecting identity-by-descent segments. iLASH scales to large, biobank-level datasets, empowering downstream analysis that uses IBD for population genetic inference and disease mapping. In addition, we demonstrate that, consistent with population genetic theory^31^, IBD is a ubiquitous component of large-scale population datasets and can provide estimates of relatedness useful in both medical and population genetic contexts. Here, we have run iLASH on large datasets such as the PAGE Study (n=51,520) and the UK Biobank (n= 487,330), providing additional context to the understanding of population structure using typical measures of global ancestry relying on unlinked genotypes. As IBD breaks down via recombination across a relatively small number of generations, the observed patterns from iLASH provide a snapshot of relatedness and population structure driven by more recent patterns, rather than the more ancient patterns determined by SNP-based genetic drift.

In contrast to previous methods, we gain significant performance improvements by basing our algorithm on locality-sensitive hashing (LSH), an algorithm that leverages the speed of hash-based methods while allowing for some incomplete matches, whether due to genotyping or phase switch-based inconsistency. By contrast, GERMLINE, the previous industry standard for large-scale inference, only identifies IBD candidates on small hash seeds, followed by SNP-wise extension, increasing runtime quadratic to the number of individuals due to a large number of pairwise comparisons. Keeping most of our algorithm within the standard LSH framework allows our runtime to grow much more slowly in the number of individuals. In addition, we include several low-level optimizations to allow for parallelizability on modern computing nodes, improving speed and sensitivity.

While this windowed hash-based method could mean that our method is less precise at identifying IBD end-points along the chromosome, in practice, our simulations show otherwise. We validated this method via simulations, ensuring that we could both recover short segments (3-5cM) as well as the full length of longer segments, allowing for downstream utility of the IBD tract length distribution. Our method is far more sensitive than GERMLINE at identifying short segments, an observation found by others^32^. Identifying large segments is critical for inferring an unbiased tract length distribution, an observation required for IBD-based genetic relatedness matrices^33^, as well as population genetic inference^3^. While maintaining high sensitivity in short segments, we ensure that our method is not susceptible to false positives via genome shuffling^25^ to create a putatively IBD-free dataset. Consistent with our method being haplotypic, false positives should not be an issue, and we observe our method being almost entirely without false positives, up to the detection limit of haplotypic complexity in our test data. We note that these false positive tests can be highly dependent on the diversity present in the population used for simulation, therefore we chose a population with limited endogamy, derived from PAGE Study African Americans, to test iLASH.

In addition to identifying cryptic relatedness in a dataset, we anticipate our scalable IBD tool to provide additional insights into large-scale datasets such as PAGE, UK Biobank, and larger datasets such as the Million Veteran Program and the All of Us Research Program. iLASH works within standard pipelines as it uses a standard input file format, and output from these estimates can easily be integrated into medical and population genetic workflows. As an example, we demonstrate the utility in estimating IBD segment patterns across real-world datasets, allowing for downstream population clustering using graphical methods able to tease apart fine-scale population structure at a level beyond standard SNP-based methods such as PCA. This, then, can have large implications for medical genetics, particularly at the rare end of the frequency spectrum, where variants are far more likely to be private to one or a small number of populations. For example, we have shown previously that IBD tracts allow us to identify a population-specific variant underlying a recessive musculoskeletal disease in Puerto Ricans, that could not be detected using standard SNP-based genome-wide association approaches^8^.

While our method is computationally efficient, mega-scale datasets in the hundreds of thousands to millions still benefit from data subsetting in the current version of iLASH, as demonstrated in running iLASH over the UK Biobank. This can be ameliorated with runs on high-memory nodes, however to fit the entirety of a dataset on a single machine will require additional data compression, likely through methods such as the Positional Burrows-Wheeler Transformation (PBWT), employed by the Sanger Imputation Service. These approaches can be integrated efficiently in the future, along with other methods, such as a distributed implementation of iLASH tailored for machine clusters, and incremental computation of IBD as new subjects are added to large datasets, such as from the UK Biobank 150,000 participants release to the current >500,000 individuals, or with direct-to-consumer companies. A distributed implementation of iLASH, designed natively to be run over nodes in a cluster, fits well with the underlying algorithm and would allow for an even more scalable solution. We have currently focused our methods on common variants as are typically found in genotype arrays. We plan in the future to update iLASH to account for recent, rare mutations as are present in sequence data. As our algorithm is based on locality-sensitive hashing we can easily handle mismatches due to genotype error or recent mutation on an IBD background. This simply will require modification of haplotype similarity thresholds and SNP density. With large, diverse sequencing datasets soon available, we anticipate this as a future improvement to a new version of iLASH.

Numerous methods have been created to model population structure for large, diverse populations. However, as datasets grow, population structure becomes more inevitable, and the relevance of demographic history influencing patterns of cryptic relatedness become unavoidable. This has particular implications for how we think of genotypic similarity. Where phasing and imputation are standard workflows, we provide a method to integrate IBD analysis into existing pipelines, allowing for novel population identification and inference of demographic history. From these we can both provide a method forward for population-based linkage as a complement to standard GWAS approaches, as well as an efficient way of identifying sub-populations within a larger dataset. Methods such as iLASH then, while having their roots firmly in early medical genetic studies, can then provide insight for the future of large-scale and multi-ethnic cohorts available in biobanks and national initiatives.

## Acknowledgments

We would like to thank Jonathan Shortt and Tonya Brunetti for initial testing. This work was supported in part by NIH grants U01HG009080, U01HG007419, R56HG010297 and R01HG010297.

## Availability

The iLASH code, manual, description of best practices, and additional scripts are available at github.com/roohy/IBD

The Population Architecture using Genomics and Epidemiology (PAGE) data is available through dbgap at accession phs000356.

## Author Contributions

J.L.A. and C.R.G. led the development of the approach. R.S. and J.L.A. designed the algorithm, with input on IBD techniques from C.R.G. R.S. implemented the algorithm and ran simulations. R.S. and G.M.B. ran the approach on the PAGE dataset. G.M.B. ran the approach on the UK Biobank dataset. R.S. and G.M.B. analyzed results, with help from E.E.K., C.R.G., and J.L.A. R.S., G.M.B., C.R.G., and J.L.A. wrote the manuscript with critical edits from E.E.K. and C.A. All authors provided revisions and edits to the submitted manuscript.

## Methods (online)

In this section, we describe in detail the algorithm and implementation techniques used in iLASH, including parameter configurations and their effect on the results of iLASH.

### Background and rationale

iLASH was inspired by a minhash-based realization of the LSH algorithm^16,17^. Locality Sensitive Hashing (LSH) refers to a category of hashing methods that preserve a specific distance function. A hash function is called “locality-sensitive” if it maps close vectors in the source domain to close or identical vectors in the target domain. A good example of such hash functions is a mapping of the points on the surface of a half-sphere to a 2D circle plane beneath them. This function reduces dimensionality from 3 to 2. However, the output of such mapping still has enough information to infer the distance among different points on the 3D curve.

LSH was developed and is often used for duplicate string detection in large text corpora. In general, it is not feasible to compare every pair of strings, since the computational cost grows quadratically with the number of strings. Moreover, it is desirable to also identify segments that are similar, but not identical, since we need to account for text modifications such as typos, local rephrasing, insertions of advertisements, personalized greetings, or other dynamically generated text in a web page. Jaccard similarity, or the intersection of two sets divided by their union, is a measure fit for such tasks.

The LSH implementation used in finding text duplicates thus tries to preserve the Jaccard similarity between different pairs of strings.^21^ The first step is to convert each string into a set of shingles (aka n-grams, substrings of n characters) and conceptually create a matrix with the strings (sets) as rows and all the distinct shingles (elements) as columns. Then, LSH estimates the Jaccard similarity between the sets (strings) by doing two levels of hashing. The first hashing step of LSH is called *minhashing*. To create the *minhash* matrix, the algorithm generates *n* random permutations of shingles. For each permutation P, it records for each set S, the index of the first shingle included in S (cf. Figure 1). The probability of two sets having the same minhash value for each of the permutations is equal to their Jaccard similarity score. The second level of hashing is called the LSH. To calculate LSH signatures, consecutive minhash values are grouped together and hashed for a second time. Suppose there are n minhash values for each string, grouped in *b* bands. Each band is comprised of *r* = *n*/*b* minhash values. Suppose S_1_ and S_2_ have a Jaccard score of *s* between them. Then the probability of all minhash values in a band being equal between the two sets is s^r^. The probability that at least one of the minhash values in a band being different is 1-s^r^. If one or more than one of the values in a band differs between S_1_ and S_2_, then the LSH signature of that band is going to be different for the two sets. Thus, the probability of all LSH signatures being distinct for each set is (1 - *s*^*r*^)^*b*^. Using this equation, we can calculate the probability of two sets sharing at least one LSH signature, leading to them being declared a hit as 1 - (1 - *s*^*r*^)^*b*^. This probability distribution is a sigmoid function that generates a S-curve with a step transition that can be manipulated by changing values of r and b to trade off reducing the number of comparisons and the false positive and false negative rates.

Parallels between finding similar strings and similar haplotypes make adopting LSH in the genomic domain attractive. However, pure LSH is not ideal to be used over entire chromosomes simply because the goal of IBD estimation is finding exact loci shared between pairs of individuals and not an average estimation of overall similarity of pairs of chromosomes. The latter is less relevant to geneticists because on average there is not a lot of difference between similarity scores of different individuals. However, when dividing genotype data into windows (we will use the word *slice* from here on to refer to these windows along the genome) along the chromosomes, the similarity score of individuals sharing an IBD tract in or around those slices would dramatically differ from that of a non-sharing pair. For example, in the problem of IBD estimation for tracts longer than 3cM in a dataset, if all the haplotypes are divided similarly into slices shorter or equal to 3cM each, three scenarios could happen to IBD segments longer than 3cM in a database.

1. The segment is located exactly within one slice’s boundaries with minimal overflow/underflow. Then the said slice will have almost 100% similarity.
2. The segment is located almost half and half between two neighboring slices. Since the segment is larger than 3cM (the length of each slice), each slice would have around 50% similarity of more.
3. The segment is spread more on one slice and less on the neighboring slice. One of the slices will have more than 50% similarity in between the two individuals and the other slice will have less than 50% shared.

In each of these scenarios, there will be at least one slice that has a Jaccard similarity score equal to or greater than 50% between the two individual haplotypes sharing IBD. iLASH estimates such similarities in order to find IBD tracts. Segments longer than 3cM have a higher minimum score between two haplotypes which is more favorable to iLASH requirements. By inspecting neighboring slices to a slice with a high degree of estimated similarity, iLASH addresses the third scenario. This will help find the true extent of an IBD segment. In the following section, we discuss how using dynamic and overlapping slicing addresses these issues.

### iLASH algorithm and settings

As discussed earlier, the iLASH algorithm has four main steps. It first starts by slicing haplotype data into slices. Initially, we used static slices of 2000 SNPs. However, genotype data is not usually sampled uniformly through the human genome. Thus, there can be slices of 2000 SNPs that cover a 5 cM long haplotype and there can be others covering 2 cM in the same experiment. To address this, we used *dynamic slicing* where each slice corresponds to a fixed genetic distance of k cM (usually 3 cM) but comprises a variable number of SNPs. For added precision, neighboring slices can overlap. For example (and across our experiments), the 3 cM slices in our experiments overlapped on 1.4 cM with their neighbors. In the areas of low complexity, where a small number of SNPs could represent a long haplotype, we observed increased rates of false-positives. Thus, we defined a threshold in iLASH to prevent the analysis of slices with lower than a given SNP count (in our experiments we ignored slices with fewer than 50 SNPs). While we have found these parameters to yield good results on our datasets, they may or may not be suitable for other datasets. Our implementation of iLASH allows the user to configure all these parameters.

Starting the second step, the SNP data in each slice is tokenized into shingles (k-mers). In our experiments, each shingle encompasses 20 SNPs. Smaller shingle length does not necessarily mean higher precision as it may cause the number of possible values for each shingle to decrease which results in lower precision. The shingles are then mapped to a 32-bit integer space using FNV hashing to allow for uniformly representing shingles of different lenghts (see Other Technical Notes below). No stored hash structure is used so as to maximize the speed gains by eliminating the need for a synchronized memory access procedure. By default, iLASH uses non-overlapping shingles. Our experiments used this default setting. However, the tool has the option to generate overlapping shingles which can help with noisy data by increasing the similarity at the cost of a modest increase in run time. Jaccard similarity of slices S_1_ and S_2_ was then defined as the number of shingles they have in common divided by the total number of shingles present in the slices. In the next two steps, iLASH calculates the minhash values and the LSH signatures. The resulting hash table comprised of LSH signatures helps iLASH algorithm to find hits between slices that are similar. The algorithm uses FNV hashing to a 32-bit integer space for both the minhashing and LSH steps. In the LSH steps, however, an in-memory hash table was maintained since synchronization is inherently critical for finding hits. In the experiments for this paper, we used 20 minhash permutations per slice. These minhash values were then grouped into 5 bands to generate LSH signatures.

In its last step, iLASH analyzes LSH hits. Getting a hit among LSH signature for two slices, does not necessarily mean the two are a match or even similar enough to be considered for further analysis. Based on the number LSH hits between two slices, iLASH estimates a minimum similarity score for them. Next, it examines the estimated similarities using two thresholds. The *Match Threshold* parameter (MT) controls iLASH decision whether to declare the two slices a match. Matching slices are not examined before being written to output. However, if an estimated score is lower than MT, it will be compared to *Interest Threshold* (IT). Scores that are not declared a match but are higher than interest threshold will be examined on a shingle level in order to find matching sub-slices. These sub-slices, if longer than the minimum length, will be written to the output file. For our experiments, we used a match threshold of 99% and interest threshold of 70%.

Finally, neighboring matched slices are collapsed together to form longer segments. Before writing them to the output file, iLASH examines the edges of each segment at a shingle level; and extends it if possible. This will help to recover as much of the actual segment as possible.

### Other Technical Notes

To maximize deployment and adoption, iLASH is designed to run on a standard single machine, without requiring parallel technologies like Hadoop or CUDA. However, iLASH takes advantage of the multiple cores available in modern machines and efficiently interleaves computation with reading and writing to disk (I/O operations). To read from and write to files, programs are required to wait in I/O queues. Furthermore, the speed of storage devices is far lower than that of main memory. Therefore, I/O operations can hinder the experiments. iLASH uses parallelization to process the genotypes that are already loaded in memory while it waits for the next batch of data to be loaded. Also, while waiting to write IBD segments to the output file, it computes the next IBD segments.

Instead of using the shingles directly, iLASH hashes the shingles using FNV hashing, which has several advantages. First, FNV hashing allows iLASH to deal with different shingle sizes uniformly. It especially helps to compress the shingle space when the shingle length parameter is set to more than 30 bits. Second, FNV hashing enables iLASH to analyze both normalized genotype files (encoded as bit strings of minor/major alleles) and unnormalized files (with for example letter encoding for the bases) in the same fashion. Third, it allows for parallelization, as opposed to using standard C++ hash tables that would create a synchronization bottleneck among the multiple threads. Finally, iLASH also uses FNV to create the permutations to compute minhash signatures. When computing minhash signatures, using FNV to map shingles to a 32-bit space and then using that representation space to create random permutations using a formula instead of actually permuting the matrix, helps maximize the effect of parallelization by eliminating the need to maintain an integrated hash table among all threads. Assuming x to be the hash value of a shingle to be the index number of that shingle, we can generate a new index number for x in a random permutation P(a,b) using the following formula; where a and b are randomly selected integers specific to the permutation:

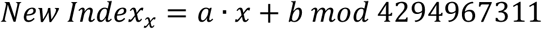

The FNV hash function is also used for generating the LSH signature. However, unlike other steps that involved hashing, analyzing LSH signatures requires maintaining a hash table in order to find hits. Removing in-memory hash structures in shingling and minhashing steps helped us to effectively parallelize our algorithm and gain a substantial speedup against our original implementation.

### Test data generation

To simulate genotype data of individuals, we took a composite individual approach^23^. Genotype data of African American individuals in the PAGE Study was broken down in short windows and randomly rearranged to eliminate IBD while preserving LD structure. We then randomly copied haplotypes between different individuals to simulate IBD for power tests.

### Application to Population Architecture using Genomics and Epidemiology (PAGE) Study data

A total of 51,520 subjects were genotyped on the MEGA array as part of the Population Architecture using Genomics and Epidemiology (PAGE) study, and subsequently underwent quality control filters as described in Wojcik *et al*.^34^. Genotypes that passed quality control (n=1,402,653) underwent phasing using SHAPEIT2. At this stage an additional N=1 individual was removed for having a chromosome specific missingess rate of > 10%. Phased haplotypes for the autosomes were subsequently filtered to a minor allele frequency of 5% and indels were removed (resulting in the retention of 593,326 autosomal SNPs genome-wide). SHAPEIT2 output was then converted to plink format using the fcgene software and these files were used as the input for both GERMLINE and iLASH (along with genetics maps interpolated using data from b37 genetic maps).

To compute IBD we used comparable parameters for GERMLINE and iLASH. The flags used for GERMLINE were *“-min_m 3 -err_hom 0 -err_het 2 -bits 25 –haploid.”*

For iLASH the parameters were:

*auto_slice 1, perm_count 12, shingle_size 20, shingle_overlap 0, bucket_count 5, perm_count 20, max_thread 20, match_threshold 0.99, interest_threshold 0.70, max_error 0, min_length 3, cm_overlap 1.4.*

The code, along with best practices and other recommendations in the online user manual, is available at https://github.com/roohy/IBD

### Quality control for downstream analysis in PAGE

IBD haplotypes inferred for N=38,919 PAGE individuals from the WHI, MEC, and HCHS/SOL studies were filtered for regions that overlapped with known genomic regions of low complexity. Additionally, IBD haplotypes that fell within genomic regions of excess IBD sharing (empirically defined as regions where we observed that the mean number of inferred IBD haplotypes exceeded 3 standard deviations of the genome-wide mean) were also excluded from downstream analysis.

### Identity-by-descent network construction and community detection

The length of IBD haplotypes (cM) shared between each pairs of PAGE individuals were summed to obtain the total length of IBD shared genome-wide between pairs. This was used as the input for the construction of an undirected network using the iGraph software in R (version 3.2.0) where each individual was represented as a node, weighted edges were used to represent the sum of IBD sharing between any give pair. Community detection was then performed using the infomap.community() function from the iGraph package.

### Application to UK Biobank Data

Quality control and phasing of the UK Biobank genotype data was performed as previously described^35^. Phased haplotype data for N=487,330 UK Biobank participants in BGEN v1.2 format were converted to vcf format using bgenix (v1.0.1) and subsequently converted to PLINK format using an in-house python script. After the removal of indels, a total of 655,532 SNPs were retained across the autosomes. These sites were used as the input for iLASH, which was run using the following parameters:

“*perm_count 12, shingle_size 20, shingle_overlap 0, bucket count 4, max_thread 20, match_threshold 0.99, intersect_threshold 0.70, max_error 0, min_length 2.9, auto_slice 1, cm_overlap 1.4”*

## References

1. Carmi, S. et al. The variance of identity-by-descent sharing in the Wright-Fisher model. Genetics 193, 911–28 (2013).

2. Erlich, Y., Shor, T., Pe’er, I. & Carmi, S. Identity inference of genomic data using long-range familial searches. Science 362, 690–694 (2018).

3. Palamara, P.F., Lencz, T., Darvasi, A. & Pe’er, I. Length distributions of identity by descent reveal fine-scale demographic history. Am J Hum Genet 91, 809–22 (2012).

4. Browning, S.R. & Browning, B.L. Accurate Non-parametric Estimation of Recent Effective Population Size from Segments of Identity by Descent. Am J Hum Genet 97, 404–18 (2015).

5. Browning, S.R. & Browning, B.L. Identity by descent between distant relatives: detection and applications. Annu Rev Genet 46, 617–33 (2012).

6. Browning, S.R. & Browning, B.L. High-resolution detection of identity by descent in unrelated individuals. Am J Hum Genet 86, 526–39 (2010).

7. Kenny, E.E. et al. Systematic haplotype analysis resolves a complex plasma plant sterol locus on the Micronesian Island of Kosrae. Proc Natl Acad Sci U S A 106, 13886–91 (2009).

8. Belbin, G.M. et al. Genetic identification of a common collagen disease in puerto ricans via identity-by-descent mapping in a health system. Elife 6(2017).

9. Kong, A. et al. Detection of sharing by descent, long-range phasing and haplotype imputation. Nat Genet 40, 1068–75 (2008).

10. O’Connell, J. et al. Haplotype estimation for biobank-scale data sets. Nat Genet 48, 817–20 (2016).

11. Loh, P.R., Palamara, P.F. & Price, A.L. Fast and accurate long-range phasing in a UK Biobank cohort. Nat Genet 48, 811–6 (2016).

12. Indyk, P. & Motwani, R. Approximate nearest neighbors: towards removing the curse of dimensionality. in Proceedings of the thirtieth annual ACM symposium on Theory of computing 604–613 (ACM, Dallas, Texas, USA, 1998).

13. Gusev, A. et al. Whole population, genome-wide mapping of hidden relatedness. Genome Res 19, 318–26 (2009).

14. Wang, J. et al. Trinary-projection trees for approximate nearest neighbor search. IEEE Trans Pattern Anal Mach Intell 36, 388–403 (2014).

15. Shrivastava, A. & Li, P. Densifying one permutation hashing via rotation for fast near neighbor search. in Proceedings of the 31st International Conference on International Conference on Machine Learning - Volume 32 I-557-I-565 (JMLR.org, Beijing, China, 2014).

16. Broder, A. On the Resemblance and Containment of Documents. in Proceedings of the Compression and Complexity of Sequences 1997 21 (IEEE Computer Society, 1997).

17. Dasgupta, A., Kumar, R. & Sarlos, T. Fast locality-sensitive hashing. in Proceedings of the 17th ACM SIGKDD international conference on Knowledge discovery and data mining 1073–1081 (ACM, San Diego, California, USA, 2011).

18. Manku, G.S., Jain, A. & Sarma, A.D. Detecting near-duplicates for web crawling. in Proceedings of the 16th international conference on World Wide Web 141–150 (ACM, Banff, Alberta, Canada, 2007).

19. Chum, O., Philbin, J., Isard, M. & Zisserman, A. Scalable near identical image and shot detection. in Proceedings of the 6th ACM international conference on Image and video retrieval 549–556 (ACM, Amsterdam, The Netherlands, 2007).

20. Henn, B.M. et al. Cryptic distant relatives are common in both isolated and cosmopolitan genetic samples. PLoS One 7, e34267 (2012).

21. Leskovec, J., Rajaraman, A. & Ullman, J.D. Mining of massive datasets / Jure Leskovec, Standford University, Anand Rajaraman, Milliways Labs, Jeffrey David Ullman, Standford University, xi, 467 pages (Cambridge University Press, Cambridge, 2014).

22. Purcell, S. et al. PLINK: a tool set for whole-genome association and population-based linkage analyses. Am J Hum Genet 81, 559–75 (2007).

23. Naseri, A., Liu, X., Zhang, S. & Zhi, D. Ultra-fast Identity by Descent Detection in Biobank-Scale Cohorts using Positional Burrows-Wheeler Transform. bioRxiv (2017).

24. Su, Z., Marchini, J. & Donnelly, P. HAPGEN2: simulation of multiple disease SNPs. Bioinformatics 27, 2304–5 (2011).

25. Fu, W., Browning, S.R., Browning, B.L. & Akey, J.M. Robust Inference of Identity by Descent from Exome-Sequencing Data. Am J Hum Genet 99, 1106–1116 (2016).

26. Durbin, R. Efficient haplotype matching and storage using the positional Burrows-Wheeler transform (PBWT). Bioinformatics 30, 1266–72 (2014).

27. Wojcik, G. et al. The PAGE Study: How Genetic Diversity Improves Our Understanding of the Architecture of Complex Traits. bioRxiv (2018).

28. Rosvall, M. & Bergstrom, C.T. Maps of random walks on complex networks reveal community structure. Proc Natl Acad Sci U S A 105, 1118–23 (2008).

29. Zhou, W. et al. Efficiently controlling for case-control imbalance and sample relatedness in large-scale genetic association studies. Nat Genet 50, 1335–1341 (2018).

30. Loh, P.R., Kichaev, G., Gazal, S., Schoech, A.P. & Price, A.L. Mixed-model association for biobank-scale datasets. Nat Genet 50, 906–908 (2018).

31. Shchur, V. & Nielsen, R. On the number of siblings and p-th cousins in a large population sample. J Math Biol (2018).

32. Bjelland, D.W., Lingala, U., Patel, P.S., Jones, M. & Keller, M.C. A fast and accurate method for detection of IBD shared haplotypes in genome-wide SNP data. Eur J Hum Genet 25, 617–624 (2017).

33. Evans, L.M. et al. Narrow-sense heritability estimation of complex traits using identity-by-descent information. Heredity (Edinb) 121, 616–630 (2018).

34. Wojcik, G.L. et al. Genetic analyses of diverse populations improves discovery for complex traits. Nature 570, 514–518 (2019).

35. Bycroft, C. et al. The UK Biobank resource with deep phenotyping and genomic data. Nature 562, 203–209 (2018).

