## Supplementary Material for "Rapid detection of identity-by-descent tracts for mega-scale datasets"

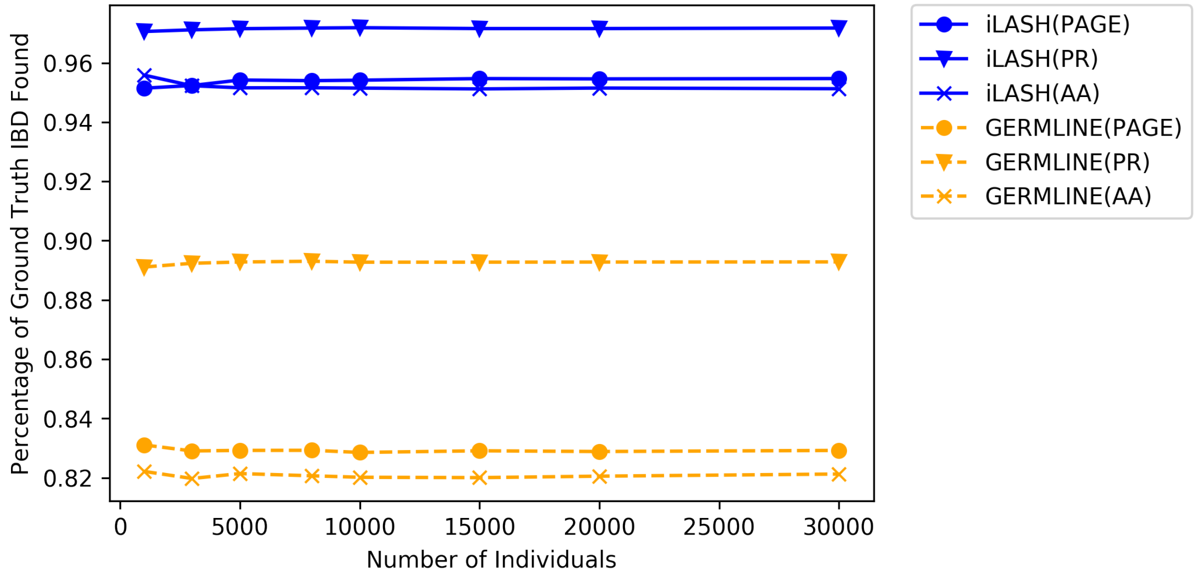


**Supplemental Figure 1.** Concordance of iLASH and GERMLINE with ground truth on simulated data from three populations based on distributions found on the PAGE study: African Americans (AA), Puerto Ricans (PR), and all subjects (PAGE).

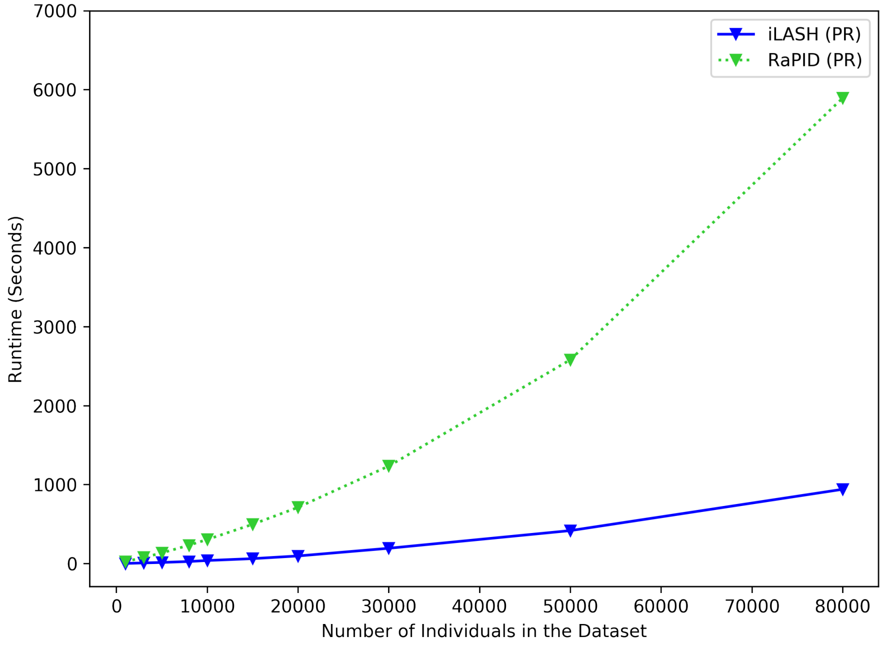


**Supplemental Figure 2.** iLASH-RaPID runtime comparison based on dataset size using simulated data derived from the Puerto Rican population in the PAGE study.

**Supplementary Table 1.**

| Community | PPV | Top Label |
|---|---|---|
| 1 | 0.954943274 | AfricanAmerican |
| 2 | 0.985674085 | NativeHawaiian |
| 3 | 0.958929226 | Mexico |
| 4 | 0.955396256 | PuertoRico |
| 5 | 0.975548061 | Japan |
| 6 | 0.978395062 | Cuba |
| 7 | 0.980787704 | DominicanRepublic |
| 8 | 0.428571429 | Mexico |
| 9 | 0.996835443 | Mexico |
| 10 | 0.972972973 | Puno |
| 11 | 0.5 | PuertoRico |
| 14 | 0.571428571 | Ecuador |
